# Bayesian estimation of population size and overlap from random subsamples

**DOI:** 10.1101/2021.07.06.451319

**Authors:** Erik K. Johnson, Daniel B. Larremore

## Abstract

Counting the number of species, items, or genes that are shared between two sets is a simple calculation when sampling is complete. However, when only partial samples are available, quantifying the overlap between two sets becomes an estimation problem. Furthermore, to calculate normalized measures of *β*-diversity, such as the Jaccard and Sorenson-Dice indices, one must also estimate the total sizes of the sets being compared. Previous efforts to address these problems have assumed knowledge of total population sizes and then used Bayesian methods to produce unbiased estimates with quantified uncertainty. Here, we address populations of unknown size and show that this produces systematically better estimates—both in terms of central estimates and quantification of uncertainty in those estimates. We further show how to use species count data to refine estimates of population size in a Bayesian joint model of populations and overlap.

## Introduction

Quantifying the overlap between two populations is a problem in many fields including genetics, ecology, and computer science. When the two populations or sets are fully known, one can simply count the size of their intersection. However, when populations are only partially observed, due to a subsampling or stochastic sampling process, the population overlap problem becomes one of inference.

In ecology, the relationship between the diversity in one population and another is called *β*-diversity [37], an idea which has led to the creation of numerous indices and coefficients which seek to quantify it. For example, the canonical Jaccard index [20] and the Sorenson-Dice coefficient [16, 32] have the appealing properties that (i) they are based only on the number of shared species, *s*, and the numbers of species in each population, *R_a_* and *R_b_*, and they take the values zero, when two populations are entirely unrelated, and one, when the populations are identical. However, these coefficients, as well as alternatives [21], have been shown to be biased when population sampling is incomplete [10, 23]. Furthermore, they provide no measure of statistical uncertainty because they provide only point estimates.

To address these issues, improvements in the quantification of *β*-diversity have been made in various ways. One direction of development recognizes that the measurement of *β*-diversity from the presence and absence of species fundamentally relies on counting the species shared by the two populations in the context of the numbers of species in each population separately, thus cataloguing the myriad ways in which these three integers might be reasonably combined, depending on the circumstances [21]. Another set of developments has been to work with species abundance data instead of binary presence-absence measurements [6]. A third set of developments has been to place observations of both abundance and presence-absence in the context of a probabilistic sampling process [10, 23], allowing for the appropriate quantification of uncertainty through confidence intervals or credible intervals.

One key feature of the *β*-diversity measures that quantify uncertainty is that the assumptions of their underlying statistical models must be stated explicitly. This provides transparency and also reveals assumptions which may not hold in practice. In recent work, a Bayesian approach to *β*-diversity estimation was introduced which provides unbiased estimates of the overlap between two stochastically sampled populations, yet this approach assumes that the two original population sizes are known a priori [23]. In practice, however, overall population sizes may be unknown, or may vary widely, making this model and others like it misspecified from the outset to an unknown degree. Thus, while incorporating appropriate uncertainty into population overlap estimation is an improvement, doing so without recognizing uncertainty or misspecification in each individual population’s size may nevertheless lead to biased, overconfident, and unreliable inferences.

Here we address this problem by leveraging an additional and often available source of data in presence-absence studies: the total number of independent samples taken from each population, i.e. the sampling depth or effort. Building on the same intuition as the estimation of total species from a species accumulation curve [17], we introduce a model for *β*-diversity calculations which produces joint estimates of *s*, *R_a_*, and *R_b_* in a Bayesian statistical framework. Posterior samples of these quantities offer solutions to issues identified above by providing unbiased central estimates, the quantification of uncertainty via credible intervals, and the construction of Bayesian versions of the canonical Jaccard and Sorenson-Dice coefficients (as well as 20 others which are based on *s*, *R_a_*, and *R_b_* [21]).

Although estimating pairwise similarity is a problem in many fields, here we present the problem in the context of estimating the genetic similarity between pairs of malaria parasites from the species *Plasmodium falciparum*—the most virulent of the human malaria parasites.

### *P. falciparum* repertoire overlap problem

Of the diverse multigene families of *P. falciparum*, the *var* family is the most heavily studied because of its direct links to both malaria’s virulence and duration of infection [2, 13, 27, 35, 25]. Each *P. falciparum* parasite genome contains a repertoire of hypervariable and mutually distinct *var* genes [18]. The *var* genes differ within and between parasites, due to rapid recombination and reassortment [14, 38]. Critically, while the number of *var* genes found in each parasite’s repertoire is typically around 60, the actual number may vary considerably [28]. For instance, the reference parasite 3D7 has been measured to have 58 *var* genes [18] while the DD2 and RAJ116 parasites have 48 and 39, respectively [30].

Recent studies of *P. falciparum* epidemiology and evolution have generated insights by comparing the *var* repertoires between parasites through *β*-diversity calculations [3, 1, 11, 4, 5, 34, 15, 12]. Indeed, since *var* repertoires are, themselves, under selection, theory suggests that if a human population has been exposed to particular *var* genes, then repertoires containing those *var* genes will have lower fitness than repertoires that are entirely unrecognized by local hosts, shaping the *var* population structure [4, 34, 7, 19, 29, 5]. Methods by which we estimate the extent to which *var* repertoires overlap are therefore important, particularly as studies of the population genetics and genetic epidemiology of *P. falciparum* antigens become more sophisticated and data rich. However, as with estimates of *β*-diversity in ecology, traditional estimates of overlap between *var* repertoires also suffer bias due to subsampling.

Due to the massive diversity and recombinant structure of *var* genes, the vast majority of *var* studies to date have been restricted to using degenerate PCR primers targeting a small “tag” sequence within a particular *var* domain called DBL*α* [33]. Although these DBL*α* tags have been widely used to study the structure and function of *var* genes [35, 3, 4, 33, 8, 9, 26, 36, 24], DBL*α* PCR data nevertheless comprise a random subsample from each parasite’s repertoire of *var* genes. Thus, these procedures produce (i) presence-absence data for various *var* types, and (ii) a count of the total number of samples accumulated in the process.

In the malaria literature, repertoire overlap, also called pairwise type sharing [3], is most commonly quantified by the the Sorenson-Dice coefficient:

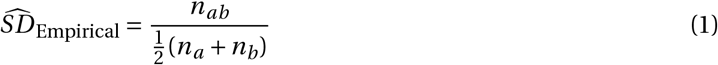

where *n_a_* and *n_b_* are the number of unique *var* types sampled from parasites *a* and *b*, respectively, and *n_ab_* is the number of sampled types shared by both parasites (i.e., the empirical overlap). When repertoires are not fully sampled (as is overwhelmingly the case in existing studies [3, 1, 11, 4, 34, 15]) the Sorensen-Dice coefficient underestimates the true overlap between repertoires. Problematically, this downward bias increases as *n_a_* and *n_b_* decrease [10, 23] resulting in difficulties when making comparisons between study sites with different sampling depths.

The methods introduced in this paper, while targeted more broadly at the development of *β*-diversity quantification, are developed in the particular context of this *P. falciparum* repertoire overlap problem.

## Methods

### Setup

Our method for inferring overlap is based on two key observations. First, not all repertoires are the same size but information about a repertoire’s size can be gleaned from the rate at which more samples identify new repertoire elements [17]. Second, the observed overlap *n_ab_* is a realization of a stochastic sampling process which depends on not only the true overlap but also the true repertoire sizes. These observations lead us to use a hierarchical Bayesian approach (Figure 1).

**Figure 1:**
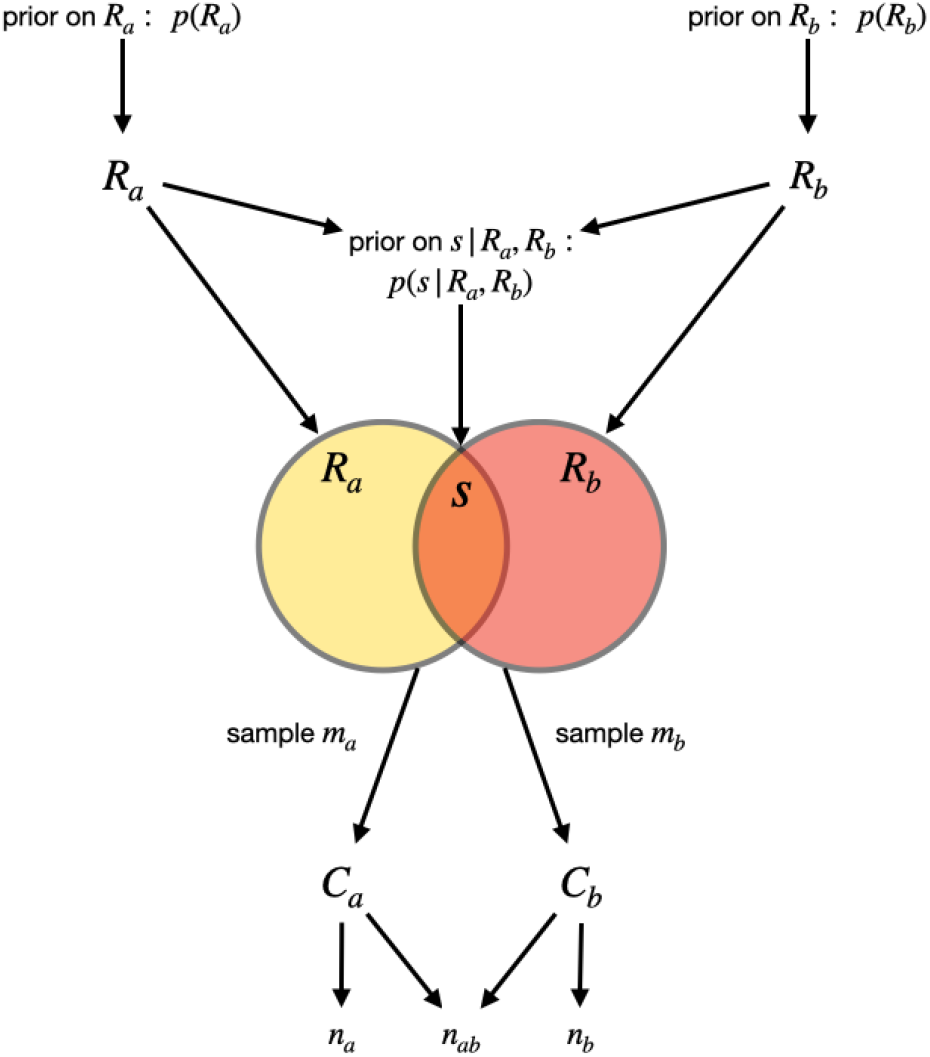
Diagram of the model. Two repertoire sizes, *Ra* and *Rb*, are generated by their priors. The overlap between the repertoires, *s*, is then generated by the prior on the overlap given the repertoire sizes. The repertoire sizes and overlap define the two parasites, *a* and *b*, from which we sample. Sampling *m a* items with replacement from parasite *a* produces count data *Ca* consisting of genes sampled from parasite *a* and counts per gene. Sampling *mb* items with replacement from parasite *b* produces count data *Cb* consisting of genes sampled from parasite *b* and counts per gene.

In brief, we model the stochastic process that generates the observed presence-absence data (*n_a_*, *n_b_*, and *n_ab_*) which can be derived from observed sample counts (i.e. observed abundances, *C_a_*, *C_b_*), from two parasites with repertoire sizes *R_a_* and *R_b_* and overlap *s*. The core of this stochastic sampling process is the assumption that sampling from each repertoire is done independently, uniformly at random, and with replacement, corresponding to PCR of *var* gDNA without substantial primer bias. From this model, we compute the joint posterior distribution of the unknown parameters, *s*, *R_a_*, and *R_b_*. With this joint posterior distribution, *p*(*s*, *R_a_*, *R_b_ | C_a_*,*C_b_*), we can produce unbiased *a posteriori* point estimates of the repertoire sizes and overlap, and can quantify uncertainty in these point estimates via credible intervals.

In the detailed methods that follow, we describe our choice of priors over the three parameters *s*, *R_a_*, and *R_b_*, derive the model likelihood, and review the steps required to make calculations efficient. An open-source implementaton of these methods is freely available (see Code Availability statement).

### Choice of prior distributions

Due to extensive sequencing and assembly efforts [28], the repertoire sizes for thousands of *P. falciparum* parasites have been characterized, leading us to choose a data-informed prior distribution for repertoire sizes *R_a_* and *R_b_*. We assume an informative Poisson prior for *R_a_* and *R_b_*, fit to the repertoire sizes from 2398 parasite isolates published by Otto et al. [28].

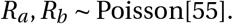

For *β*-diversity studies outside of *P. falciparum*, alternative informative priors can be chosen. Because the repertoire overlap *s* can take values between 0 and min{*R_a_*, *R_b_*}, we use an uninformative prior for repertoire overlap *s*,

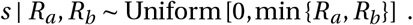

### Computing the joint posterior distribution *p*(*s*, *R_a_*, *R_b_ | C_a_*,*C_b_*)

The posterior distribution of the parameters given the count data is a product of three terms

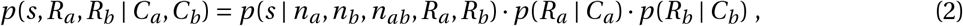

a calculation shown in detail in Appendix 1. The rest of this section is devoted to computing each of these terms, noting that that last two are mathematically identical, but derived from different data.

To compute *p*(*R | C*), the distribution of repertoire size given count data for a fixed but arbitrary total sampling effort *m*, we first calculate the likelihood of observing count data *C* given a repertoire size *R*, i.e., *p*(*C | R*). Knowing how to compute *p*(*C | R*), allows us to calculate *p*(*R | C*) via Bayes’ rule

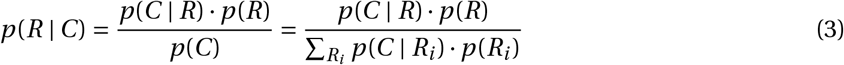

where *p*(*R*) is the prior on repertoire size and the sum in the denominator should be computed over the support of *p*(*R*). For the unbounded support of the Poisson prior used here, we restrict the sum to only those terms above the numerical precision of the computer.

In Appendix 2, we prove that

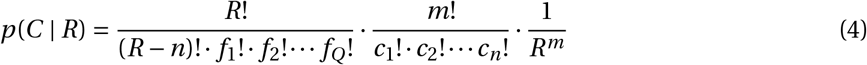

where the *c_i_* are the number of times each of the *n* sampled *var* types were observed and the *f_i_* are the multiplicities of the unique numbers in 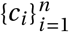. For instance, suppose the count data consists of five unique *var* types with counts

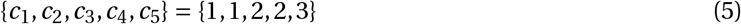

then there are three (*Q =* 3) unique numbers amongst the *c_i_* : 1, 2, and 3. Further, 1’s multiplicity in {1, 1, 2, 2, 3} is 2, 2’s is 2, and 3’s is 1 so ( *f*_1_, *f*_2_, *f*_3_) = (2, 2, 1).

With the likelihood *p*(*C | R*) in hand, it is straightforward to calculate the posterior *p*(*R | C*) via Equation (3). And, thus, we can calculate the second and third terms in Equation (2).

Conveniently, the remaining term of Eq. (2) *p*(*s | n_a_*, *n_b_*, *n_ab_*, *R_a_*, *R_b_*) has been derived in the literature [23], but only under the restriction that *R_a_* = *R_b_* = 60. We therefore rederive this quantity for general but fixed *R_a_* and *R_b_*, summarizing the main steps here.

Using Bayes’ rule, we can write

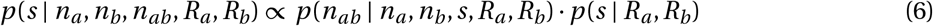

where *p*(*s R_a_*, *R_b_*) is a user-specified prior described above. The other term, *p* (*n_ab_* | *n_a_*,*n_b_*, *s*, *R_a_*,*R_b_*), can be computed by considering the probability that two subsets of size *n_a_* and *n_b_* will have an intersection of size *n_ab_*, given that they have been drawn uniformly from sets of total size *R_a_* and *R_b_* whose intersection is size *s*. To do so, we use the hypergeometric distribution, 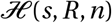, which is the distribution of the number of “special” objects drawn after *n* uniform draws with replacement from a set of *R* objects, *s* of which are “special.” With this distribution in mind, note that observing *n_ab_* shared *var* genes can be thought of as a two-step process. First, draw *n_a_ var* genes from parasite *a*’s *R_a_* total in which *s* are special because they are shared with parasite *b*. The number of shared *var*s drawn is a random variable 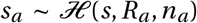. Second, draw *n_b_* genes from parasite *b*’s *R_b_* total in which *s_a_* are special because they are shared by both parasites *and* were drawn from parasite *a*. The number of shared *var*s captured after sampling from both parasites, *n_ab_*, will be distributed according to 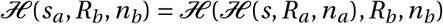.

To generate a particular empirical overlap *n_ab_*, first step 1 must happen and then, independently, step 2 must happen. We therefore multiply these two hypergeometric probabilities. However, because these two steps may occur for any value of the intermediate variable *s_a_*, we sum over all possible values of *s_a_*

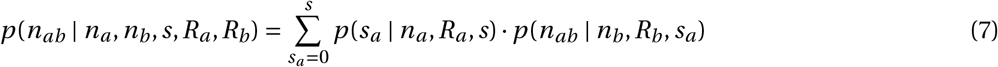

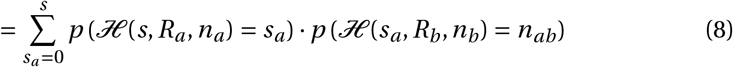

Plugging this into Equation (6) allows us to compute *p*(*s | n_a_*, *n_b_*, *n_ab_*, *R_a_*, *R_b_*).

### Inference Method Summary

We now have all the pieces in place to compute *p*(*s*, *R_a_*, *R_b_ | C_a_*,*C_b_*):

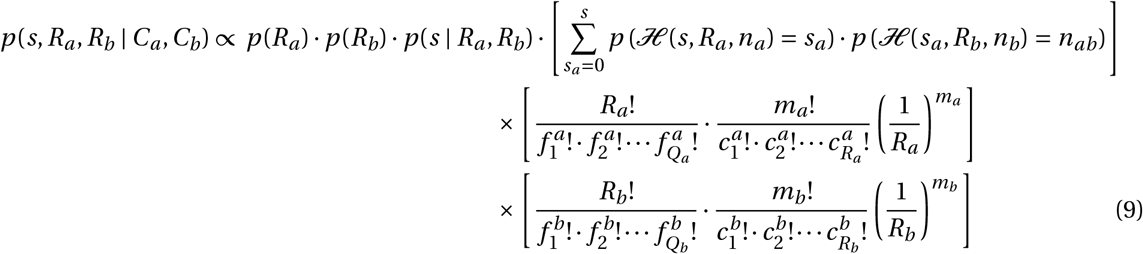

where the first three terms are the user-specified priors. With this joint posterior distribution, we can compute unbiased Bayesian estimates of *s*, *R_a_*, and *R_b_* as expectations over the posterior:

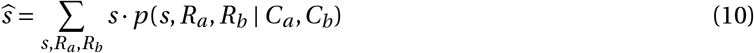

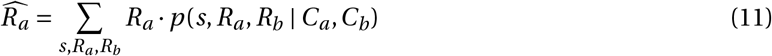

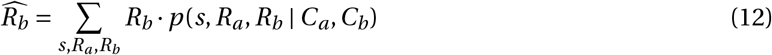

Moreover, and importantly, we can compute unbiased Bayesian estimates of any functional combination of *s*, *R_a_*, and *R_b_* such as Bayesian versions of the Jaccard index [20], the Sorensen-Dice coefficient [32], other coefficients based on *s*, *R_a_*, and *R_b_* [21], and the directional pairwise-type-sharing measures of He et al. [19]. For all of these measures, in addition to the point estimates, the ability to draw from the joint posterior distribution Eq. (9) enables one to compute credible intervals to quantify uncertainty.

### Generation of Simulated Data

To facilitate numerical experiments in which we tested our inference method’s ability to recover accurate estimates of *s*, *R_a_*, and *R_b_*, we generated synthetic data via simulation as follows. First, we selected a value of overlap *s* between 0 and 70, so that analyses could be stratified according to overlap. Next, we drew repertoire sizes *R_a_* and *R_b_* independently from the prior distribution, ensuring that *R_a_ ≤ s* and *R_b_ ≤ s*, redrawing as necessary. Next, we drew from the model (Fig. 1) a set of *m_a_* and *m_b_* samples from repertoires of sizes *R_a_* and *R_b_*, respectively, with specified overlap *s*, to generate count data histograms *C_a_* and *C_b_*. This procedure therefore stochastically created synthetic count data for a specified overlap *s* and sampling depth *m*, allowing us to test our method’s accuracy and uncertainty quantification under various scenarios.

## Results

### Inference

We first investigated how increasing the total number of independent samples *m* improves our ability to correctly estimate the total population size *R*. To do so, we conducted numerical experiments where we presumed a repertoire size and then simulated samples from it to produce count data *C*. An example of such an experiment shows how posterior estimates approach the true repertoire size as sampling effort *m* increases (Fig. 2). The true value of *R* is always contained within the inferred distributions, but only when the number of samples *m* grows large are inferences about *R* highly confident.

**Figure 2:**
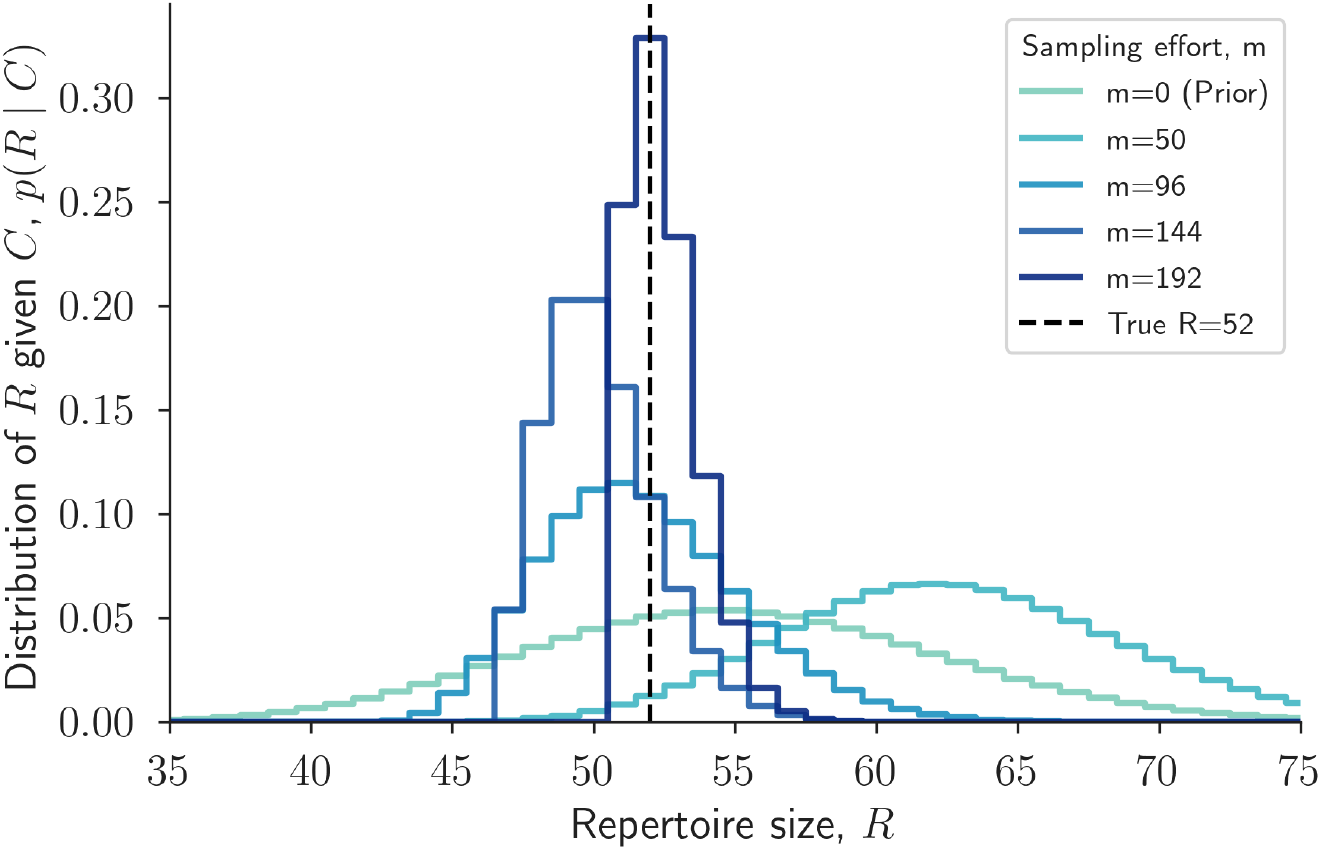
Repertoire size posterior estimates improve with increased sampling effort. For true repertoire size *R =* 52, the posterior distribution *p*(*R | C*) is plotted for different sampling efforts *m* (see legend). For each value of *m*, count data *C* were generated by drawing *m* genes uniformly with replacement from a repertoire of 52 genes. As sampling effort increases, the posterior *p*(*R | C*) concentrates around the true repertoire size 52. The *m =* 0 curve is the Poisson prior on repertoire size, *p*(*R*).

This experiment illustrates two related points. First, there is valuable information in knowing the total sampling effort *m*, even if some samples were duplicate observations of previously observed genes, simply because those sample counts inform repertoire size estimates. Second, increasing the sampling effort concentrates *p*(*R | C*) around the true repertoire size, concretely linking sampling effort to estimation of not only repertoire size, but through decreased uncertainty, eventual overlap estimates as well.

Next we examined whether the *s*, 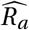, and 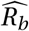 estimates in Equations (10)–(12) are accurate across a range of sampling efforts *m* in two steps. First, we simulated the sampling process for various values of *s*, *R_a_*, and *R_b_* to produce synthetic count data *C_a_* and *C_b_* with varying levels of overlap between the observed samples. Then, we evaluated our ability to recover *s*, *R_a_*, and *R_b_* by applying Eqs. (10)–(12) to the synthetic data.

We found that the overlap and repertoire estimates accurately reproduce the true parameter values, provided that sampling effort is sufficiently large. Furthermore, as sampling effort increases, estimates become increasingly accurate (Fig. 3).

**Figure 3:**
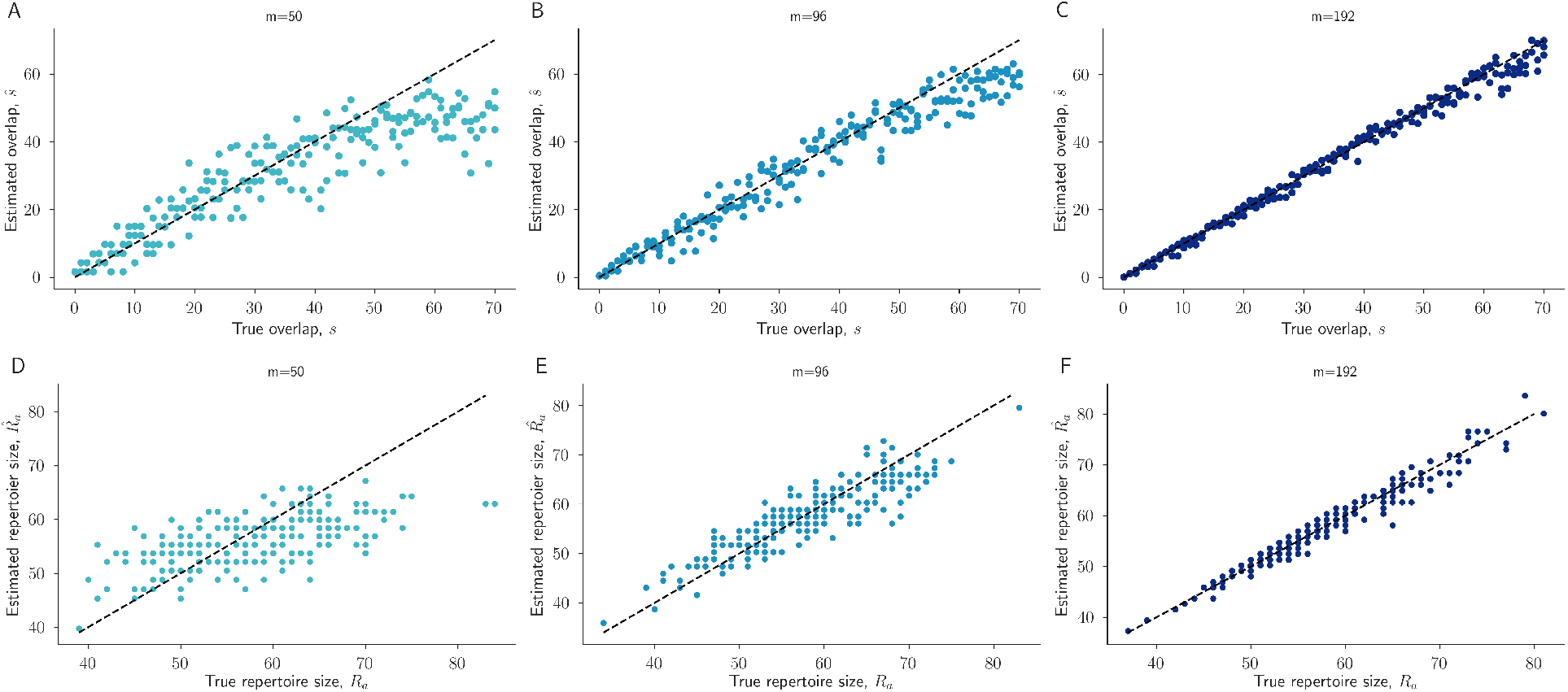
Accuracy of estimates across a range of true parameter values and sampling efforts. For each overlap value *s* between 0 and 70, we performed three independent simulations to generate synthetic count data (Methods).. Estimates of *s* (A,B,C) and *R_a_* (D,E,F) from the resulting count data, using our statistical model, are shown. Estimates are shown for sampling efforts *m* 50, 96, 192 across left, middle, and right columns, respectively. Dashed black lines represent perfect unbiased inference.

However, we also observed that when the sampling effort is small but repertoires are large and highly overlapping (e.g. *m =* 50 and *s >* 50), *ŝ* underestimates the true values (Fig. 3A). This phenomenon is due to a more general property of Bayesian inference: when there are fewer samples from which to infer, the prior distribution exerts a stronger effect on inferences. Here, the Poisson prior over repertoire sizes assigns low probability to repertoire sizes as large as 70 (*p*(*R_a_* ≥ 70) = 0.03), and thus, in the absence of a large sampling effort to overwhelm that prior, the surprisingly large repertoire sizes and overlaps require substantially more samples *m* to establish. In real data from *P. falciparum*, repertoires (and thus repertoire overlaps) larger than 60 are rarely observed [28, 15], decreasing the potential impact of this issue.

### Uncertainty

Bayesian methods also allow us to quantify uncertainty via credible intervals (CIs). To measure how well our CIs capture the true parameter values, we computed 95% highest density posterior intervals for parameter estimates in simulated data, where true values were known. As expected, uncertainty decreased as sampling effort increased, and approximately 95% of the 95% CIs captured the true parameter values, as designed (Fig. 4). For instance, for sampling efforts of *m* 50, *m* 96, and *m* = 192, the proportions of the 95% *ŝ* CIs containing the true *s* were 0.975, 0.975, and 0.965, respectively. For the same three sampling efforts, the proportions of the 95% 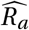 CIs that contained the true repertoire size *R_a_* were 0.920, 0.950, and 0.955, respectively.

**Figure 4:**
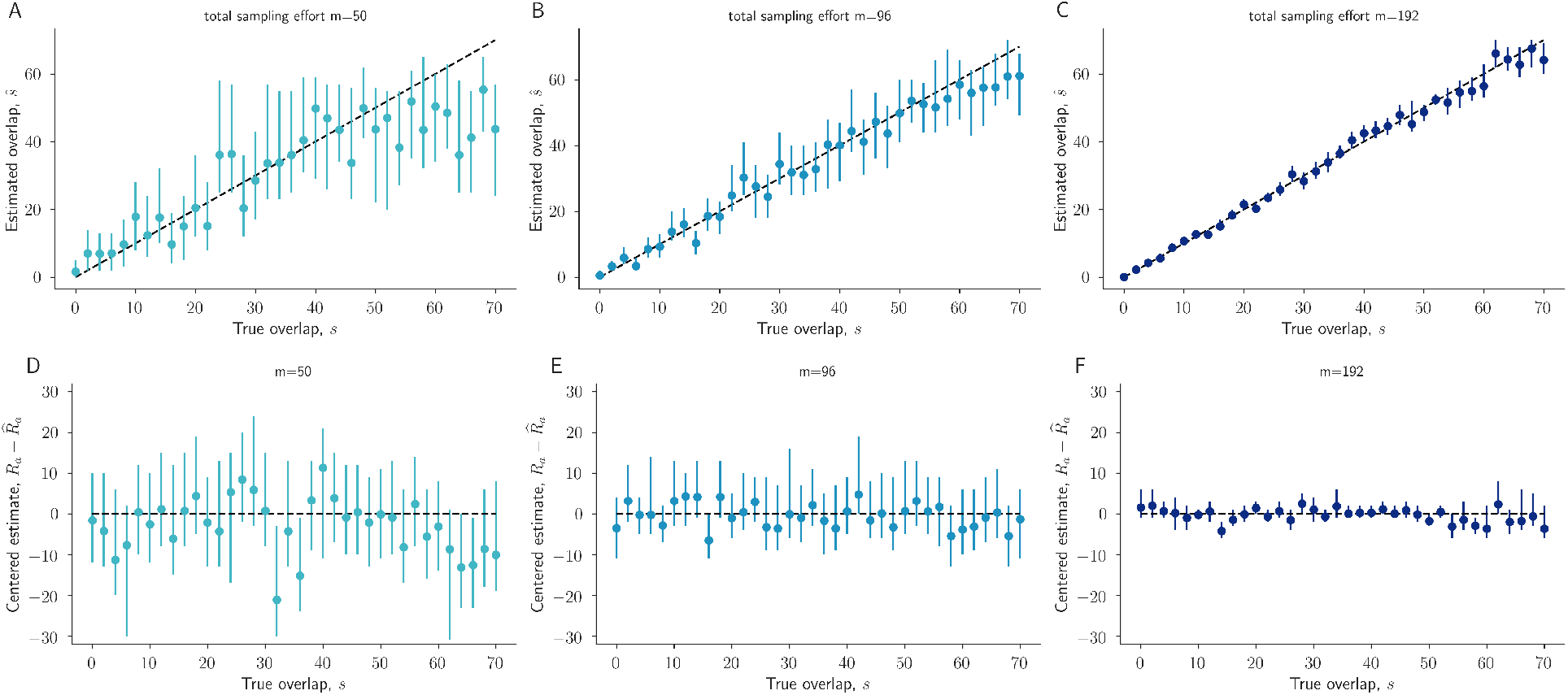
Credible intervals quantify uncertainty in overlap estimates. For each overlap value *s* between 0 and 70, we performed one simulation to generate synthetic count data (Methods). Estimates from the resulting count data, using our statistical model, of *s* (A,B,C), and error in *R_a_* and *R_b_* (D,E,F) are shown. Estimates (dots) and 95% credible intervals (lines) are shown for sampling efforts *m* 50, 96, 192 in left, middle, and right columns, respectively.

### Improving *β*-diversity indices

Over 20 different indices of *β* diversity have been proposed which algebraically combine empirical estimates of *R_a_*, *R_b_*, and *s* [21], including the well known Jaccard index and the Sorenson-Dice coefficient. The Sorenson-Dice coefficient is defined as the ratio of repertoire overlap to the average of the repertoires sizes,

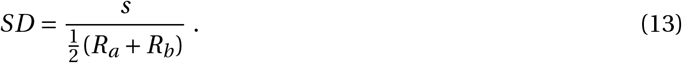

Typically, in the absence of more sophisticated estimates of *R_a_*, *R_b_*, and *s*, empirical values are used,

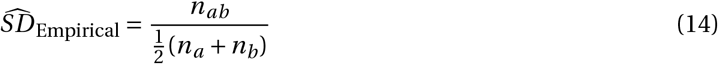

However, the joint posterior distribution Eq. (9) over *s*, *R_a_*, and *R_b_* opens the door to a Bayesian reformulation of the Sorenson-Dice coefficient as

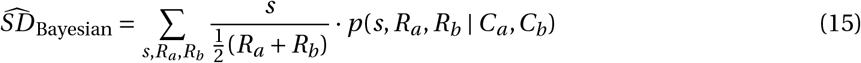

with similar generalizations for the Jaccard coefficient or other combinations of *s*, *R_a_*, and *R _b_* [21]. This Bayesian Sorenson-Dice coefficient a verages the values of the typical Sorenson-Dice coefficient over joint posterior estimates of *s*, *R_a_*, and *R_b_*.

We investigated the performance of the Bayesian Sorenson-Dice coefficient 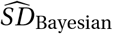 and its empirical counterpart 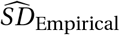 by once more simulating the sampling process under known conditions and applying both formulas. As in our estimates of repertoire overlap, we again found that Bayesian Sorenson-Dice estimates produce consistent and unbiased estimates with correct quantification of uncertainty via credible intervals (Fig. 5), except when sampling effort is low (*m =* 50) while true repertoire overlap is extremely high (*s* > 50). Furthermore, the Bayesian estimates track the true Sorenson-Dice values better than direct empirical estimates across overlap values and sampling efforts; direct empirical estimates are biased more and more downward as sampling effort decreases and as true overlap increases (Fig. 5). While this illustrates how the Bayesian framework herein may be used to improve classical and commonly used estimators via Eq. (15), an identical approach may be used to compute Bayesian Jaccard coefficients, or other algebraic combinations of *s*, *R_a_*, and *R_b_* [21].

**Figure 5:**
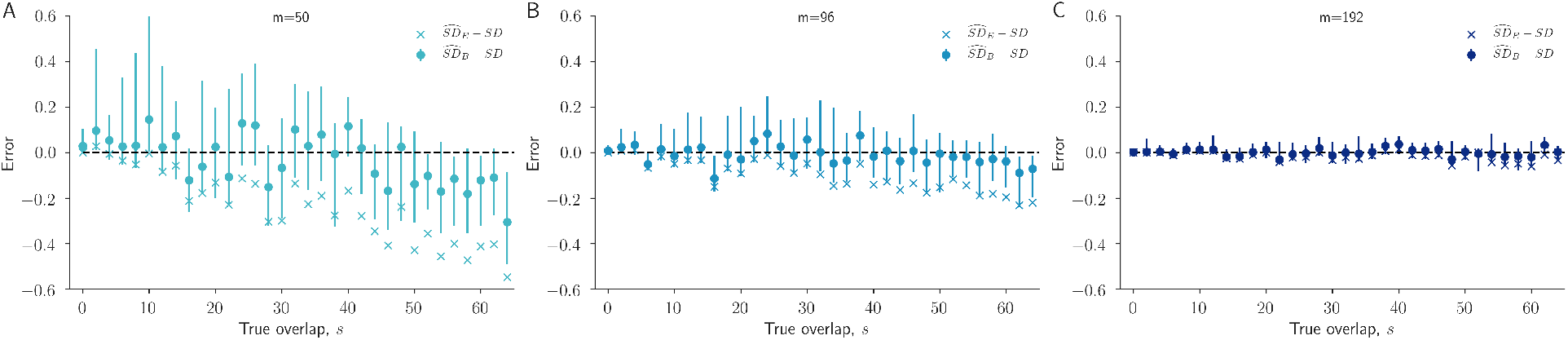
Bayesian vs empirical Sorensen-Dice estimates. For each overlap value *s* between 0 and 70, we performed one independent simulation to generate synthetic count data (Methods) and estimated the Sorensen-Dice coefficient using estimates from our Bayesian framework as well as from the raw empirical data. The error in the Bayesian Sorensen-Dice estimate, 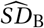 (Equation (15)), and accompanying 95% credible intervals are shown. The often-used empirical Sorensen-Dice estimate, 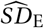 (Equation (14)), is also shown. The dashed black line at 0 represents the true Sorensen-Dice coefficient (Equation (13)).

### Sample size calculations

Sample size calculations ask how many samples are needed to produce eventual estimates with a pre-specified level of (or upper bound on) statistical uncertainty. Such questions, while critical in the ethical study of human subjects, are also important when budgeting for studies in which additional samples require time, reagents, and funding.

To assist in sample size calculations, we used simulations to quantify the relationship between increases in sampling effort *m* and decreases in the typical width of the credible interval around the repertoire overlap estimate estimate *ŝ* (Eq. (10)). For many overlap-sampling effort pairs, (*s*, *m*), we performed 300 independent replicates in which we generated synthetic data, computed the posterior distribution for *s*, and calculated the width of the 95% *ŝ* CI.

We found that, as expected, increased sampling effort *m* leads to decreased uncertainty across all values of overlap *s* (Fig. 6). However, we also found that overlap plays a role as well, with larger overlap causing wider CIs. For instance, after *m =* 200 samples, a CI for overlap *s =* 70 is typically of width 8, while a CI for overlap *s =* 30 is typically of width 4. After *m =* 300 samples from each repertoire, median CI widths are 4 or lower for all overlap values. In short, it is easier to show with high confidence that two samples do not overlap than to show that they are highly overlapping.

**Figure 6:**
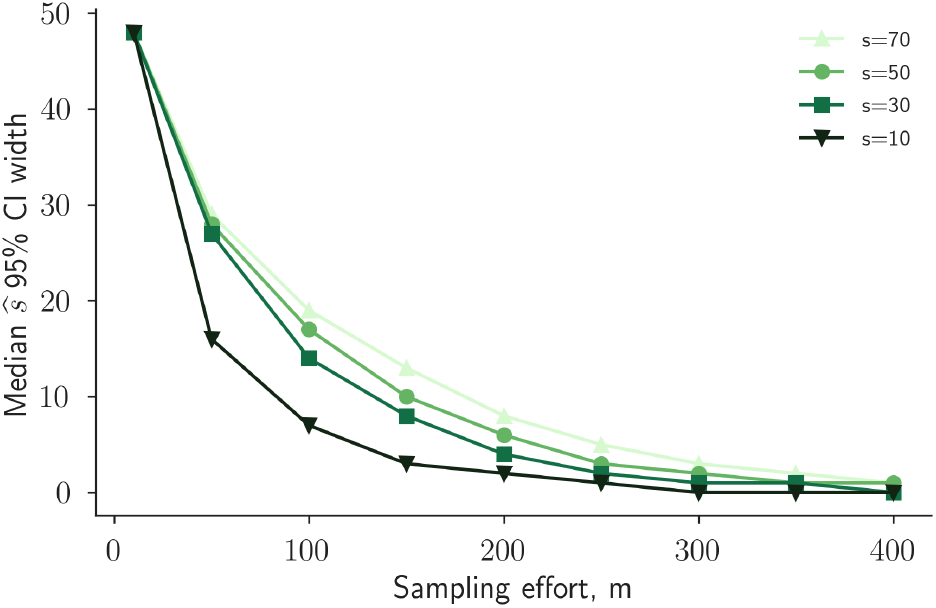
Quantifying the decrease in uncertainty from increased sequencing. Constant *s* curves show the median 95% credible interval (CI) width for the *s* estimate, *ŝ*, as a function of the sampling effort *m*. For each (*s*, *m*)-duplet, the median is across 300 count data generation simulations. This plot illustrates the intuition that additional laboratory efforts (increasing *m*) lead to higher accuracy (smaller CIs).

## Discussion

This manuscript presents a Bayesian solution to estimating the overlap between two populations when only subsamples of those populations are available. Importantly, because the total population sizes bear on the inference of overlap, this method jointly estimates population sizes and overlap from the quantitative accumulation of evidence, improving inferences. Samples from the joint posterior distribution can be used to quantify uncertainty via credible intervals, or can be used in Bayesian versions of the Jaccard index, Sorenson-Dice coefficient, and other algebraic combinations of set sizes and intersections.

In addition to the analysis of existing data, this approach can also be used prospectively to perform sample size calculations. Importantly, context-specific sample sizes can be estimated by including additional information in the Bayesian prior. For instance, in the context of malaria’s *var* genes, it is known that parasites from South America tend to have smaller repertoires [22, 31] than samples from other regions [28]—information which can be expressed through the prior distribution to influence (and in this case, decrease) sampling needs. Because additional sampling has financial and complexity costs, this allows researchers to weigh accuracy requirements against laboratory costs in the contexts of a particular study.

Beyond the study of *P. falciparum*, the approach introduced in this work lands in between two existing classes of *β*-diversity measures in the ecology literature. One class of methods measures *β*-diversity in terms of species presence or absence [21], while the other further includes species abundance [10]. The present work uses abundance measurements (which we call count data) in order to improve presence-absence-based *β*-diversity estimates, but does not construct abundance-based similarity measures per se. By drawing inferences both from this work also aligns with past efforts which rely in principle on an idea that one may draw inferences both from what is observed and what is not observed [10, 23].

The tradeoffs for improved inferences are twofold. First, our approach requires abundance data (i.e., count data *C*) instead of presence/absence totals *n_a_*, *n_b_*, and *n_ab_*. This limits the retrospective analysis of past work or meta-analyses to only those studies that meet a greater data-sharing burden. However, we also note that, as proven in Appendix 2, full count data are not necessary: the posterior *p*(*s*, *R_a_*, *R_b_ | C_a_*,*C_b_*) can still be computed exactly when only the sampling efforts (*m_a_* and *m_b_*) and the presence/absence values (*n_a_*, *n_b_*, and *n_ab_*) are known.

The second tradeoff for improved inference is that one must specify a prior distribution for the total population sizes. In the case of the *var* gene repertoires of *P. falciparum*, data-informed prior distributions can be created for both global [28] or local [31] estimates. In this light, one may view past work on Bayesian methods for repertoire overlap [23, 5] as specifying point priors at a particular fixed repertoire size. In general, the choice of an appropriate prior is left to the user, which may require users to make explicit their prior beliefs about population size.

There are limitations to our approach which relate to our assumptions about the sampling process which generates the count data. Specifically, we have assumed throughout this work that each time a new sample is generated, this sample is drawn independently and uniformly from a population in which unique genes, species, or objects are identically represented. Thus, unlike abundance based measures [10] which assume that some species are more likely to be sampled than others, we assumed each species’ selection is equiprobable.In the sampling of *var* gene sequences, for instance, methodological artifacts such as PCR primer bias may cause non-uniform sampling. One avenue for future work could be to extend our rigorous probabilistic modeling to the non-uniform sampling regime.

## Acknowledgements

The authors wish to thank Shazia Ruybal-Pesantez, Kathryn Tiedje, Karen Day, and Thomas Otto for the generosity of their feedback. This work was supported in part by the SeroNet program of the National Cancer Institute (1U01CA261277-01).

## Code Availability

All code needed to evaluate the conclusions in the paper are present in the paper and/or the Supplementary Materials, and open-source code is available^1^. The Bayesian models were implemented in Python 3.8.

## Ethics Declaration

E.K.J. and D.B.L. declare no competing interests.

## Appendix 1 Factorization of the joint posterior distribution

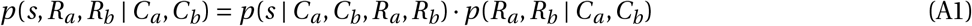

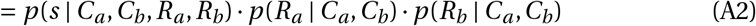

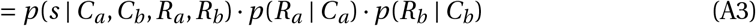

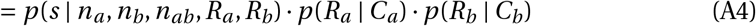

The first equality is an application of the probability identity *p*(*A*, *B*) = *p* (*A* | *B*)*p*(*B*). The second equality uses the independence of *R_a_* and *R_b_*. For the third equality, note that the count data for parasite *b* contains no pertinent information relative to parasite *a*’s repertoire size that parasite *a*’s own count data does not contain. Thus, *p*(*R_a_* | *C_a_*,*C_b_*) = *p*(*R_a_* | *C_a_*) and, similarly, *p*(*R_b_ | C_a_*,*C_b_*) = *p*(*R_b_* | *C_b_*). The fourth equality is the claim that

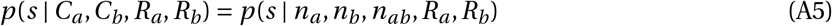

which follows from the fact that the number of times each gene was observed (i.e., the counts) informs the repertoire size as the example above showed. However, when the repertoire sizes are known, only the *n_a_*, *n_b_*, and *n_ab_* values from the count data are pertinent to the overlap size.

## Appendix 2 Theorems enabling efficient computations

## Theorem 1

*Let C be the count data resulting from sampling m elements uniformly with replacement from a set with R elements. Let n be the number of unique elements drawn from C. Then, when R is known, we can think of C as a vector*

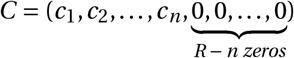

*where the c_i_ correspond to the number of times each sampled element was drawn and we have 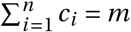. Let u = (u_1_, u_2_, …, u_Q_) be the unique **nonzero** numbers in C and let f_i_ be the number of times u_i_ appears in C. Then, the distribution of C | R is given by*

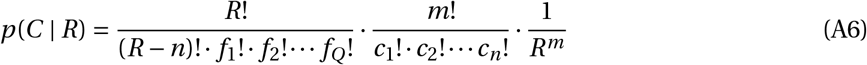

## Proof

First note that, given *R labeled* elements each with equal probability of being sampled, the multinomial distribution gives the probability of observing any given count data. This is almost the probability that we are interested in except that, for us, the elements are not labeled. That is, as an example, count data *C* (2, 3) is the same as *C* = (3, 2). So, *p*(*C* | *R*) is the multinomial probability multiplied by the number of unique permutations of the counts. The multinomial probability is given by

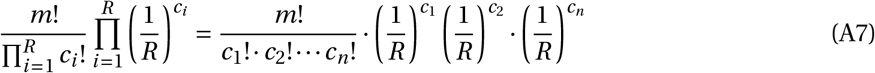

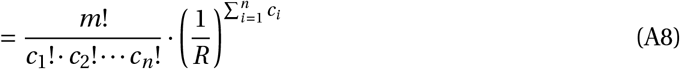

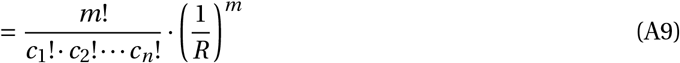

The number of unique permutations of the counts is the same as the number of *R*-letter words containing *Q* + 1 unique letters, *u*_0_, *u*_1_, …, *u_Q_*, where letter *u_i_* appears *f_i_* times for *i* ≠ 0 and letter *u*_0_ appears *R* − *n* times. This number is given by the multinomial coefficient

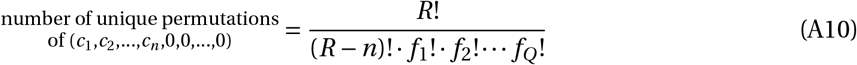

And, thus,

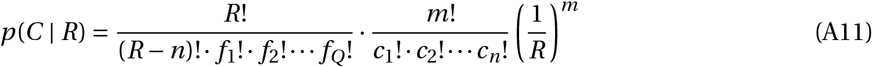

## Theorem 2.

*Let C be the count data resulting from sampling m elements uniformly with replacement from a set with R elements. The count data C consists of the unique elements sampled and the number of times each element was sampled. Let n be the number of unique elements sampled and let p(R) be the prior distribution on the (unknown) set size R. Then, for fixed C and m,*

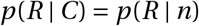

That is, *p*(*R* | *C*) depends only on the unique number of elements sampled n and not the number of times each element was sampled.

## Proof

First note it is impossible for the set size *R* to be less than number of unique elements sampled *n*. So, when *R* < *n*, *p*(*R* | *C*) = 0.

For *R* ≥ *n*, we can think of the fixed count data *C* as a vector

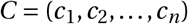

where *n* is the number of elements sampled and the *c_i_* are the number of times each sampled element was sampled. From Theorem 1, we know that

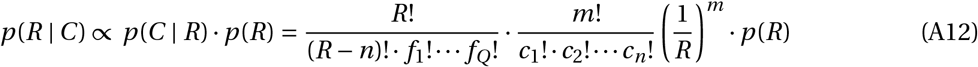

where *u*_1_, *u*_2_, …, *u_Q_* are the unique numbers *other than zero* in *C* and *f_i_* is the number of times *u_i_* appears in *C*. Dropping all the terms that do not depend on *R* gives

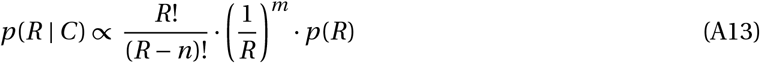

Now let’s look at *p*(*R* | *n*). First, using Bayes’ rule and ignoring the denominator term that does not depend on *R*, we have

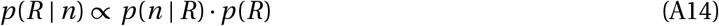

*p*(*n* | *R*) is the probability that *n* unique elements were sampled from a set with *R* elements after *m* uniform draws with replacement. To draw *n* elements after *m* draws, note that draw a previously unseen element must have been drawn *n* times and already seen elements must have been drawn *m* − *n* times. We can think of this process as a Markov chain with *R* + 1 states corresponding to the number of unique elements drawn. For the Markov chain’s probability transition matrix, note that if *i* unique elements have already been drawn then the probability that the next element drawn has already been drawn is *i* /*R* and the probability that it is a previously unseen element is (*R* − *i*)/*R*. Thus, the probability transition matrix *π* has entries

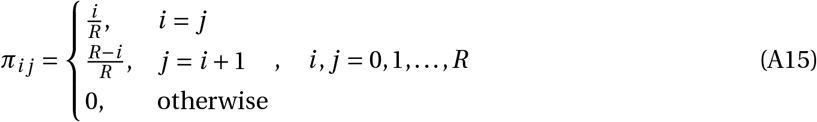

To calculate *p*(*n* | *R*), we will sum over all possible paths that the Markov chain could have taken to get from state 0 to state *n* in *m* steps. Let’s denote all the possible paths that start at state 0 and end at state *n* after *m* steps by 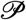. Since every possible path must have must start at 0 and end at *n*, every possible path must include the following transitions: 0 → 1, 1 → 2, …, and *n* − 1 → *n*. The remaining *m* − *n* steps must have been steps for which the number of unique elements drawn did not change, i.e., a previously drawn element was drawn again. So the possible paths are differentiated by the number of times *q_i_* that the chain stayed in state *i*. For notational convenience, let *Q* be the set of all unique *n*-tuples (*q*_1_, *q*_2_, …, *q_n_*) such that each *q_i_* is a nonnegative integer and 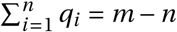. In this notation, summing over paths is equivalent to summing over the *n*-tuples in *Q*

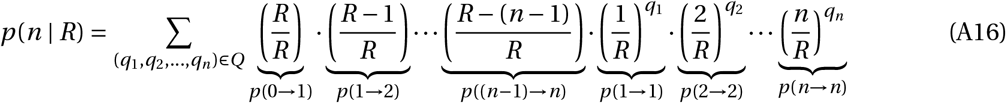

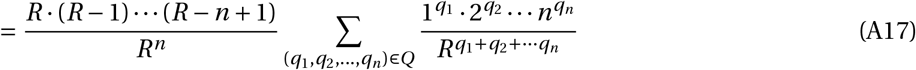

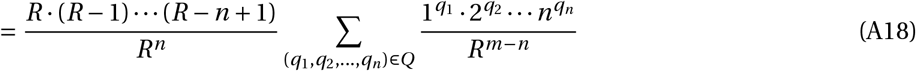

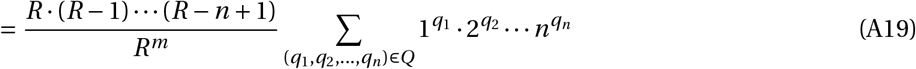

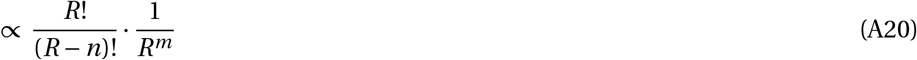

where, in the last equation, we have dropped the sum that doesn’t depend on *R* and used the fact that 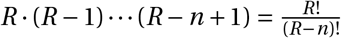

Plugging this result into *p*(*R* | *n*) gives

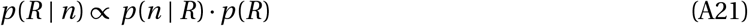

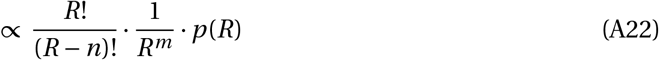

which, as a function of *R*, is the same expression we found for *p*(*R* | *C*). Thus, for fixed count data *C*,

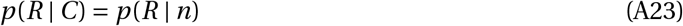

In the context of estimating *var* repertoire sizes and assuming PCR samples *var*s uniformly, this result means that only knowing the sampling effort *m* and the number of unique *var*s sampled *n* is as informative as knowing all the counts per gene.

## Footnotes

1 https://github.com/erikj540/Bayesian-Beta-Diversity

## References

[1] Letusa Albrecht, Catarina Castineiras, Bruna O Carvalho, Simone Ladeia-Andrade, Natal Santos da Silva, Erika HE Hoffmann, Rosimeire C dalla Martha, Fabio TM Costa, and Gerhard Wunderlich. The South American *Plasmodium falciparum var* gene repertoire is limited, highly shared and possibly lacks several antigenic types. Gene, 453(1-2):37–44, 2010.

[2] Marion Avril, Abhai K Tripathi, Andrew J Brazier, Cheryl Andisi, Joel H Janes, Vijaya L Soma, David J Sullivan, Peter C Bull, Monique F Stins, and Joseph D Smith. A restricted subset of *var* genes mediates adherence of *Plasmodium falciparum*-infected erythrocytes to brain endothelial cells. Proceedings of the National Academy of Sciences, 109(26):E1782–E1790, 2012.

[3] Alyssa E Barry, Aleksandra Leliwa-Sytek, Livingston Tavul, Heather Imrie, Florence Migot-Nabias, Stuart M Brown, Gilean AV McVean, and Karen P Day. Population genomics of the immune evasion (*var*) genes of *Plasmodium falciparum*. PLOS Pathogens, 3(3):e34, 2007.

[4] Amy K Bei, Ababacar Diouf, Kazutoyo Miura, Daniel B Larremore, Ulf Ribacke, Gregory Tullo, Eli L Moss, Daniel E Neafsey, Rachel F Daniels, Amir E Zeituni, et al. Immune characterization of *Plasmodium falciparum* parasites with a shared genetic signature in a region of decreasing transmission. Infection and Immunity, 83(1):276–285, 2015.

[5] Amy K Bei, Daniel B Larremore, Kazutoyo Miura, Ababacar Diouf, Nicholas K Baro, Rachel F Daniels, Allison Griggs, Eli L Moss, Daniel E Neafsey, Awa B Deme, et al. Plasmodium falciparum population genetic complexity influences transcriptional profile and immune recognition of highly related genotypic clusters. bioRxiv, 2020.

[6] J Roger Bray and John T Curtis. An ordination of the upland forest communities of southern wisconsin. Ecological monographs, 27(4):326–349, 1957.

[7] Caroline O Buckee, Peter C Bull, and Sunetra Gupta. Inferring malaria parasite population structure from serological networks. Proceedings of the Royal Society B: Biological Sciences, 276(1656):477–485, 2009.

[8] Peter C Bull, Matthew Berriman, Sue Kyes, Michael A Quail, Neil Hall, Moses M Kortok, Kevin Marsh, and Chris I Newbold. *Plasmodium falciparum* variant surface antigen expression patterns during malaria. PLOS Pathogens, 1(3):e26, 2005.

[9] Peter C Bull, Sue Kyes, Caroline O Buckee, Jacqui Montgomery, Moses M Kortok, Chris I Newbold, and Kevin Marsh. An approach to classifying sequence tags sampled from *Plasmodium falciparum var* genes. Molecular and Biochemical Parasitology, 154(1):98, 2007.

[10] Anne Chao, Robin L Chazdon, Robert K Colwell, and Tsung-Jen Shen. A new statistical approach for assessing similarity of species composition with incidence and abundance data. Ecology Letters, 8(2):148–159, 2005.

[11] Donald S Chen, Alyssa E Barry, Aleksandra Leliwa-Sytek, Terry-Ann Smith, Ingrid Peterson, Stuart M Brown, Florence Migot-Nabias, Philippe Deloron, Moses M Kortok, Kevin Marsh, et al. A molecular epidemiological study of *var* gene diversity to characterize the reservoir of *Plasmodium falciparum* in humans in Africa. PLOS One, 6(2):e16629, 2011.

[12] Lauren M Childs and Daniel B Larremore. Network models for malaria: Antigens, dynamics, and evolution over space and time. 2021.

[13] Antoine Claessens, Yvonne Adams, Ashfaq Ghumra, Gabriella Lindergard, Caitlin C Buchan, Cheryl Andisi, Peter C Bull, Sachel Mok, Archna P Gupta, Christian W Wang, et al. A subset of group A-like *var* genes encodes the malaria parasite ligands for binding to human brain endothelial cells. Proceedings of the National Academy of Sciences, 109(26):E1772–E1781, 2012.

[14] Antoine Claessens, William L Hamilton, Mihir Kekre, Thomas D Otto, Adnan Faizullabhoy, Julian C Rayner, and Dominic Kwiatkowski. Generation of antigenic diversity in plasmodium falciparum by structured rearrangement of var genes during mitosis. PLoS genetics, 10(12):e1004812, 2014.

[15] Karen P Day, Yael Artzy-Randrup, Kathryn E Tiedje, Virginie Rougeron, Donald S Chen, Thomas S Rask, Mary M Rorick, Florence Migot-Nabias, Philippe Deloron, Adrian JF Luty, et al. Evidence of strain structure in *Plasmodium falciparum var* gene repertoires in children from Gabon, West Africa. Proceedings of the National Academy of Sciences, 114(20):E4103–E4111, 2017.

[16] Lee R Dice. Measures of the amount of ecologic association between species. Ecology, 26(3):297–302, 1945.

[17] Ronald A Fisher, A Steven Corbet, and Carrington B Williams. The relation between the number of species and the number of individuals in a random sample of an animal population. The Journal of Animal Ecology, pages 42–58, 1943.

[18] Malcolm J Gardner, Neil Hall, Eula Fung, Owen White, Matthew Berriman, Richard W Hyman, Jane M Carlton, Arnab Pain, Karen E Nelson, Sharen Bowman, et al. Genome sequence of the human malaria parasite *Plasmodium falciparum*. Nature, 419(6906):498–511, 2002.

[19] Qixin He, Shai Pilosof, Kathryn E Tiedje, Shazia Ruybal-Pesántez, Yael Artzy-Randrup, Edward B Baskerville, Karen P Day, and Mercedes Pascual. Networks of genetic similarity reveal non-neutral processes shape strain structure in *Plasmodium falciparum*. Nature Communications, 9(1):1–12, 2018.

[20] Paul Jaccard. Étude comparative de la distribution florale dans une portion des alpes et des jura. Bull Soc Vaudoise Sci Nat, 37:547–579, 1901.

[21] Patricia Koleff, Kevin J Gaston, and Jack J Lennon. Measuring beta diversity for presence–absence data. Journal of Animal Ecology, 72(3):367–382, 2003.

[22] Susan M Kraemer, Sue A Kyes, Gautam Aggarwal, Amy L Springer, Siri O Nelson, Zoe Christodoulou, Leia M Smith, Wendy Wang, Emily Levin, Christopher I Newbold, et al. Patterns of gene recombination shape var gene repertoires in plasmodium falciparum: comparisons of geographically diverse isolates. BMC genomics, 8(1):1–18, 2007.

[23] Daniel B Larremore. Bayes-optimal estimation of overlap between populations of fixed size. PLOS Computational Biology, 15(3):e1006898, 2019.

[24] Daniel B Larremore, Sesh A Sundararaman, Weimin Liu, William R Proto, Aaron Clauset, Dorothy E Loy, Sheri Speede, Lindsey J Plenderleith, Paul M Sharp, Beatrice H Hahn, et al. Ape parasite origins of human malaria virulence genes. Nature communications, 6(1):1–11, 2015.

[25] Thomas Lavstsen, Louise Turner, Fredy Saguti, Pamela Magistrado, Thomas S Rask, Jakob S Jespersen, Christian W Wang, Sanne S Berger, Vito Baraka, Andrea M Marquard, et al. *Plasmodium falciparum* erythrocyte membrane protein 1 domain cassettes 8 and 13 are associated with severe malaria in children. Proceedings of the National Academy of Sciences, 109(26):E1791–E1800, 2012.

[26] Johan Normark, Daniel Nilsson, Ulf Ribacke, Gerhard Winter, Kirsten Moll, Craig E Wheelock, Justus Bayarugaba, Fred Kironde, Thomas G Egwang, Qijun Chen, et al. PfEMP1-DBL1*α* amino acid motifs in severe disease states of *Plasmodium falciparum* malaria. Proceedings of the National Academy of Sciences, 104(40):15835–15840, 2007.

[27] Lucy B Ochola, Bethsheba R Siddondo, Harold Ocholla, Siana Nkya, Eva N Kimani, Thomas N Williams, Johnstone O Makale, Anne Liljander, Britta C Urban, Pete C Bull, et al. Specific receptor usage in *Plasmodium falciparum* cytoadherence is associated with disease outcome. PLOS One, 6(3):e14741, 2011.

[28] Thomas D Otto, Sammy A Assefa, Ulrike Böhme, Mandy J Sanders, Dominic Kwiatkowski, et al. Evolutionary analysis of the most polymorphic gene family in *falciparum* malaria. Wellcome Open Research, 4, 2019.

[29] Shai Pilosof, Qixin He, Kathryn E Tiedje, Shazia Ruybal-Pesántez, Karen P Day, and Mercedes Pascual. Competition for hosts modulates vast antigenic diversity to generate persistent strain structure in plasmodium falciparum. PLoS biology, 17(6):e3000336, 2019.

[30] Thomas S Rask, Daniel A Hansen, Thor G Theander, Anders Gorm Pedersen, and Thomas Lavstsen. Plasmodium falciparum erythrocyte membrane protein 1 diversity in seven genomes–divide and conquer. PLoS computational biology, 6(9):e1000933, 2010.

[31] Shazia Ruybal-Pesántez, Fabián E Sáenz, Samantha Deed, Erik K Johnson, Daniel B Larremore, Claudia Vera-Arias, Kathryn E Tiedje, and Karen P Day. Clinical malaria incidence following an out-break in ecuador was predominantly associated with plasmodium falciparum with recombinant variant antigen gene repertoires. medRxiv, 2021.

[32] Th A Sorensen. A method of establishing groups of equal amplitude in plant sociology based on similarity of species content and its application to analyses of the vegetation on Danish commons. Biol. Skar., 5:1–34, 1948.

[33] Helen M Taylor, Susan A Kyes, Christopher I Newbold, et al. *Var* gene diversity in *Plasmodium falciparum* is generated by frequent recombination events. Molecular and Biochemical Parasitology, 110(2):391–397, 2000.

[34] Sofonias K Tessema, Stephanie L Monk, Mark B Schultz, Livingstone Tavul, John C Reeder, Peter M Siba, Ivo Mueller, and Alyssa E Barry. Phylogeography of *var* gene repertoires reveals fine-scale geospatial clustering of *Plasmodium falciparum* populations in a highly endemic area. Molecular Ecology, 24(2):484–497, 2015.

[35] George M Warimwe, Gregory Fegan, Jennifer N Musyoki, Charles RJC Newton, Michael Opiyo, George Githinji, Cheryl Andisi, Francis Menza, Barnes Kitsao, Kevin Marsh, et al. Prognostic indicators of life-threatening malaria are associated with distinct parasite variant antigen profiles. Science Translational Medicine, 4(129):129ra45–129ra45, 2012.

[36] George M Warimwe, Thomas M Keane, Gregory Fegan, Jennifer N Musyoki, Charles RJC Newton, Arnab Pain, Matthew Berriman, Kevin Marsh, and Peter C Bull. *Plasmodium falciparum var* gene expression is modified by host immunity. Proceedings of the National Academy of Sciences, 106(51):21801–21806, 2009.

[37] Robert Harding Whittaker. Vegetation of the Siskiyou mountains, Oregon and California. Ecological Monographs, 30(3):279–338, 1960.

[38] Xu Zhang, Noah Alexander, Irina Leonardi, Christopher Mason, Laura A Kirkman, and Kirk W Deitsch. Rapid antigen diversification through mitotic recombination in the human malaria parasite plasmodium falciparum. PLoS biology, 17(5):e3020191, 2019.

